# Coronavirus activates a stem cell-mediated defense mechanism that reactivates dormant tuberculosis: implications in COVID-19 pandemic

**DOI:** 10.1101/2020.05.06.077883

**Authors:** Lekhika Pathak, Sukanya Gayan, Bidisha Pal, Joyeeta Talukdar, Seema Bhuyan, Sorra Sandhya, Herman Yeger, Debabrat Baishya, Bikul Das

## Abstract

We postulate that similar to bacteria, adult stem cells may also exhibit an innate defense mechanism to protect their niche. Here, we provide preliminary data on stem cell based innate defense against a mouse model of coronavirus, murine hepatitis virus-1 (MHV-1) infection. In a mouse model of mesenchymal stem cell (MSC) mediated *Mycobacterium tuberculosis* (*Mtb*) dormancy, MHV-1 infection in the lung exhibited 20 fold lower viral loads than the healthy control mice, suggesting the potential enhancement of an anti-MHV-1 defense by *Mtb*. This defense mechanism involves the *in vivo* expansion and reprogramming of CD271+MSCs in the lung to an enhanced stemness phenotype. The reprogrammed MSCs facilitate the activation of stemness genes, intracellular Mtb replication, and extracellular release of *Mtb*. The conditioned media of the reprogrammed MSCs exhibit direct anti-viral activity in an *in vitro* model of MHV-1 induced toxicity to type II alveolar epithelial cells. Thus, our data suggest that reprogrammed MSCs exert a unique innate defense against MHV-1 by activating dormant *Mtb.* The molecular details of this anti-viral defense mechanism against coronavirus could be further studied to develop a vaccine against COVID-19.

## INTRODUCTION

The severe acute respiratory syndrome coronavirus 2 (SARS-CoV-2) mediated coronavirus disease 2019 (COVID-19) pandemic demonstrates the ability of an emerging virus to create chaos in our modern health care system and a severe strain on the global economics. In the long term, post-pandemic, the SARS-CoV-2 might activate dormant bacterial infections. As per prior history, tuberculosis is one of the key bacterial infections affected by viral pandemics (*1*–*5*). Strikingly, one quarter of the world population is already infected with dormant tuberculosis (TB) (https://www.who.int/news-room/fact-sheets/detail/tuberculosis). If SARS-CoV-2 infects these dormant TB populations, it may cause severe impact in global health and economics by causing both COVID-19 and dormant TB reactivation. Thus, there is an urgent need to study the association of COVID-19 with dormant TB reactivation to avoid a later global TB pandemic.

Viral infection such as the influenza virus or SARS-CoV-1 is known to cause transient immune suppression that leads to reactivation of dormant bacterial infection (*6*–*7*,*4*). In 1918, the Spanish flu pandemic led to the rise of pulmonary TB incidence (*2*,*8*). Influenza in patients with TB had the highest death rate (*8*–*9*). In the year 2009 during the influenza A (H1N1) pandemic, a worse prognosis of influenza was observed in patients with TB or MDR/TB (*10*–*11*). Interestingly, SARS-CoV-1 and Middle East respiratory syndrome coronavirus (MERS-CoV) infected patients were reported to develop pulmonary TB (PTB) (*5*,*4*). Using a mouse model, an earlier study showed that influenza A virus cause rapid development of PTB lesions (*12*) and an increase in the mycobacterium load in the liver (*13*). Another mouse model of Influenza A virus and *Mtb* co-infection led to enhanced *Mtb* growth by a type-I IFN signaling pathway (*6*). However, severe inflammation in the lung is a common outcome of coronavirus infection (*14*), a symptom which is also commonly observed in TB patients (*15*). Thus, it is possible that coronavirus infection causing inflammation may also lead to reactivate dormant *Mtb* in the lung, which has not yet been studied.

We have been investigating TB dormancy in the adult stem cell niches (*16*–*18*). These stem cells reside in the bone marrow (BM) niche (*19*) and in the area of inflammation (*20*). We have identified a rare fraction of cells in the BM, the CD271+BM-mesenchymal stem cells (CD271+BM-MSCs) as the potential niche for dormant *Mtb* in mice and in successfully treated TB patients (*17*). In this stem cell niche, *Mtb* remains dormant, maintaining reactivation potential. Importantly, we have developed a mouse model of stem cell mediated *Mtb* dormancy (*17*). Briefly, streptomycin dependent mutant 18b strain infected mice exhibit lung infection following 3 weeks of streptomycin treatment. These mice develop granulomas, and also acquire humoral immunity against the bacteria. Following streptomycin starvation for 6 months, the bacteria acquire a non-replicating status. Mostly, these bacteria can be detected in the CD271+MSCs of BM, but a few are also present in the CD271+MSCs of lung (*17*). Furthermore, *Mtb* harboring CD271+MSCs reside in the hypoxic niche of BM (18). Notably, in this model, the presence of *Mtb*-CFUs in the non-CD271+MSC compartment of lung can be utilized as a sign of *Mtb* reactivation. Additionally in this model, the potential existence of altruistic stem cell (ASC) defense mechanism can also be studied by evaluating the enhanced stemness phenotype of CD271+MSCs as previously described (*21*–*22*,*18*).

Using this mouse model of stem cell mediated *Mtb* dormancy we sought to find out, if coronavirus can reactivate dormant *Mtb* (d*Mtb*). We are using a mouse coronavirus strain, the murine hepatitis virus -1 (MHV-1) that represents clinically relevant human coronavirus SARS-CoV-1 infection (*23*–*25*). MHV-1 infection causes acute lung inflammation by inducing acute respiratory infection (ARI) within 2-4 days in C57BL/6 mice by increasing viral load. Animals exhibit an elevated level of pro-inflammatory cytokines such as TNF-alpha during 2-14 days post-infection (*23*,*26*) and then fully recover (*23*). Therefore, this MHV-1 infected C57BL/6 mouse could be utilized for *Mtb* reactivation study.

Here, we demonstrate that MHV-1 infection causes d*Mtb* reactivation in the mouse model of stem cell mediated *Mtb* dormancy. Furthermore, we found that MHV-1 reprogram the CD271+MSCs to an enhanced stemness phenotype. This phenotype then exert altruistic stem cell (ASC) defense and this is evidenced by cytoprotection against MHV-1 infected type II alveolar epithelial (ATII) cells. Thus, the remarkable significance of this study is that although, MHV-1 infection causes d*Mtb* reactivation, it also activates an ASC based innate defense mechanism against the virus and that could be further explored to develop therapeutics to target coronavirus.

## RESULTS

### MHV-1 infection leads to reactivation of d*Mtb* in C57BL/6 mice

We reasoned that MHV-1 infection increases the *Mtb*-CFU level in the mice lungs containing dMtb harboring MSC, and thus, could reactivate pulmonary TB. To test this hypothesis, MHV-1 (5×10^3 PFU) was intranasally administered to C57BL/6 mice (n=25) (*23*,*25*), having lung CD271+ MSCs, that contain streptomycin-dependent mutant *Mtb* strain of 18b in non-replicating dormant state for 6 months (*17*,*27*). Simultaneously, mice were also injected with streptomycin as the mutant 18b strain requires the antibiotic to replicate (*27*). Study was ended after three weeks, because (i) three weeks of streptomycin is sufficient to establish lung infection (*17*), (ii) two weeks post MHV-1 infection, the C57BL/6 mice recover from viral pneumonitis (*23*,*25*), which ethically permit us to extend the study to third week. Thus, the study was ended on 20^th^ day following MHV-1 infection. During 20^th^ days of MHV-1 infection period, animals (n=5) were sacrificed at stipulated time, lungs obtained, and the viral titers as well as *Mtb*-CFUs in the lung tissues were evaluated (Figure 1B-C). We also measured the TNF-alpha in the animals’ blood on day-2 of MHV-1 infection to confirm MHV-1 induced immune response (*23*). First, we evaluated the viral infection in the mice as shown in Figure 1B. MHV-1 infected healthy mice were used as the control. In this control group, streptomycin was also added to find out if addition of streptomycin can affect the viral load. MHV-1 infection in the control group led to an increase in viral titers in the mouse lung for the first 4 days, and then gradually decreased within next two weeks, which is consistent with the previous result of C57BL/6 mice (*25*). MHV-1 infected mice with d*Mtb* (henceforth, d*Mtb*MHV-1 group) showed an increase in viral load for the first 4 days similar to control, but then rapidly subsided at the end of two-weeks, where the viral load was 20-fold less than control group (p<0.004; Figure 1B). The TNF-alpha levels in both the groups were increased by 3-4 fold (p<0.03; n=4) on day-2 (*dMtb*–32.5 ±11.4 vs7.3±4.5pg/ml; *dMtb*MHV-1-36.5 ±14.7 vs9.5 ± 3.6pg/ml). These data suggest that MHV-1 infection activated viral replication, and associated immune response in both groups, although, viral load decreased in *Mtb*-infected versus non-infected mice.

**Figure1:**
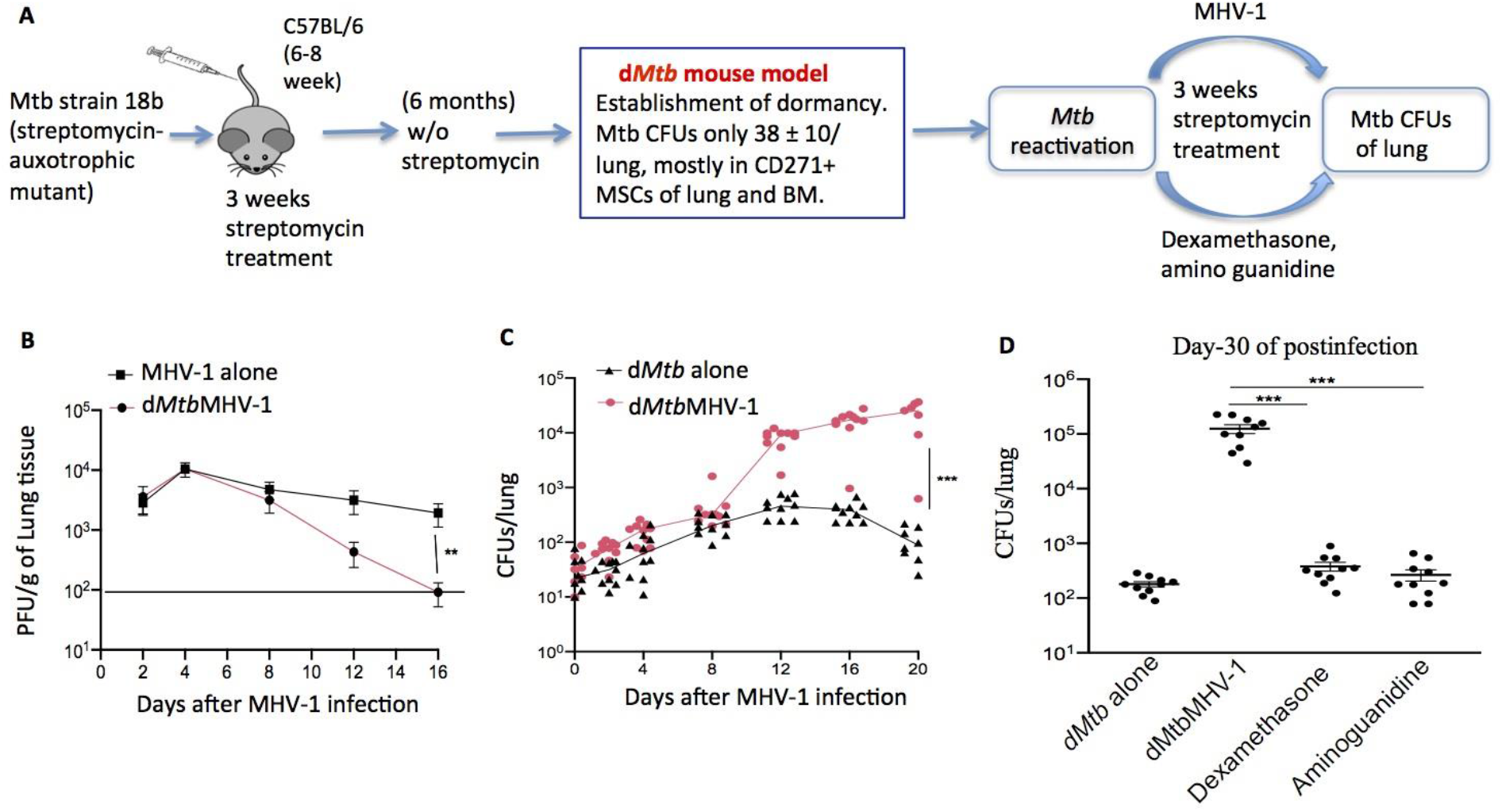
MHV-1 infection reactivates dormant Mtb in the lungs. **A.** Schematic of producing the dormant Mtb (dMtb) mouse model, and the experimental design for dMtb reactivation. **B&C.** Viral load (plaque forming unit-PFU) and Mtb-colony forming units (Mtb-CFUs) in the lungs from MHV-1 infected C57BL/6 mice at various times post infection. Solid line in B represents limit of detection. **D**. Mtb-CFUs in the lungs of mice from day-30 post MHV-1 infection, or day-30 post treatment with dexamethasone or aminoguanidine. All the mice were treated with daily streptomycin six-days per week for three weeks. **P< 0.001, ***P<0.0001, Student’s t-test; Error bar represents SEM; Mtb-CFU/lung, n= 4 independent experiments. MHV-1 alone = healthy adult mice infected with MHV-1 only. The dMtb alone = mice harboring dormant Mtb (dMtb) intracellular to lung and BM CD271+ MSCs. dMtbMHV-1 = dMtb harboring mice infected with MHV-1.

Second, we evaluated *Mtb*-CFUs in the lungs of d*Mtb* alone versus d*Mtb*MHV-1 group. Both the groups (n=10 mice in each group) received daily streptomycin treatment for mutant strain to replicate. By 8 days of infection *Mtb*-CFUs was 6-7 fold greater than the *Mtb*-CFU prior to the start of the infection in both the groups (p<0.0001, Figure 1C). Then, between days-8 and 20 of post-infection, *Mtb*-CFUs increased by 110-fold (p<0.0001) in the d*Mtb*MHV-1 group but, decreased by 2.1-fold (p<0.004) in the d*Mtb* alone group. These results indicate that MHV-1 infection not only initiated but also sustained the *Mtb* reactivation process following streptomycin treatment. In contrast, *Mtb* reactivation did not occur in the d*Mtb* alone group as the streptomycin treatment alone failed to increase *Mtb*-CFUs even after three weeks of treatment. Importantly, we found 630 fold increases in Mtb-CFU between days 0-20 in the dMtbMHV-1 group, which is equivalent to 1000 fold increase in Mtb-CFU in a dexamethasone and aminoguanidine induced Mtb reactivation model (*28*).

Therefore, next we compared the MHV-1 induced reactivation of the d*Mtb* with dexamethasone, or aminoguanidine. These have been found to reactivate d*Mtb* in a drug-induced Cornell model by suppressing the immune system (*28*). To compare the reactivation of MHV-1 versus these immunosuppressive agents, d*Mtb* harboring mice were treated with either of these two drugs and streptomycin daily for a month, then the lung *Mtb*-CFUs were quantified. Next, the *Mtb*-CFU of these two drugs treated mice with the *Mtb*-CFU obtained from one-month post-MHV-1 infected mice were compared. We found that *Mtb*-CFUs increased by only 3-4-fold (p=0.06; n=4) following either dexamethasone or aminoguanidine treatment, which was 400-fold lower than the MHV-1 infection group (p<0.00001, n=4; Figure 1D). Thus, MHV-1 infection was superior to dexamethasone or aminoguanidine to induce reactivation in our model of d*Mtb*. Our results also indicate that mechanisms other than immunosuppression may underlie MHV-1 infection mediated disease reactivation.

### MHV-1 infection causes expansion of lung CD271+MSCs and extracellular pathogen release

In this mouse model of *Mtb* dormancy, d*Mtb* remains dormant in CD271+MSCs of bone marrow and lung. We hypothesized that during d*Mtb* reactivation, the bacteria transfer from MSCs to non-MSC compartments in the lung (*17*). Thus, we analyzed the lung CD271+MSCs for intracellular *Mtb* replication and release into the extracellular space. Lung CD271+MSCs were obtained by immunomagnetic sorting after a stipulated time period of MHV-1 infection, Viable cell count was obtained by trypan blue to evaluate expansion of these cells. The d*Mtb* group served as a control. We also studied possible CD271+MSCs expansion in healthy mice infected with MHV-1 to evaluate a potential effect of this virus on MSCs of lung. Next, the sorted cells were cultured in serum-free media for 8 hours to evaluate both the intracellular and the extracellular release of *Mtb* by measuring *Mtb-*CFUs in the cell pellet and supernatant respectively. We found that the d*Mtb*MHV-1 group showed a transient 12-fold (p<0.001) expansion of CD271+MSCs between day-0 and 8. Strikingly, we also found that MHV-1 infection alone group exhibited expansion of CD271+MSCs.

However, the expansion was 6-fold lower than the d*Mtb*MHV-1 group; the d*Mtb* alone group showed no expansion of the MSCs (Figure 2B). Notably, in the d*Mtb*MHV-1 group, along with MSC expansion, there was a corresponding 27-fold (p<0.0001) increase in the number of intracellular *Mtb*-CFUs (Figure 2C). During this period supernatant of dMtbMHV-1 group did not exhibit marked increase in extracellular *Mtb*-CFUs compared to d*Mtb* control (Figure 2D). However, in the next four days i.e. between day-8 and 12, supernatant showed a 40-fold increase of *Mtb*-CFU (p<0.0032; Figure 2D). Thus, *Mtb*-transfer from CD271+ MSCs to non-CD271+MSC compartment probably occurred between the day-8 and 12, when the supernatant showed a sharp increase in *Mtb*-CFUs. Indeed, there was a corresponding 15-fold rise of *Mtb*-CFUs in the non-CD271+MSC compartment of lung cells *in vivo* (p<0.0001; Figure 2E). As expected, the d*Mtb* group showed no significant evidence of CD271+ MSC expansion, intracellular *Mtb* replication, and their extracellular release (Figure 2B-E). Altogether, the results suggest that MHV-1 infection induces a transient expansion of *Mtb* harboring CD271+MSCs and pathogen’s release into the extracellular space.

**Figure 2:**
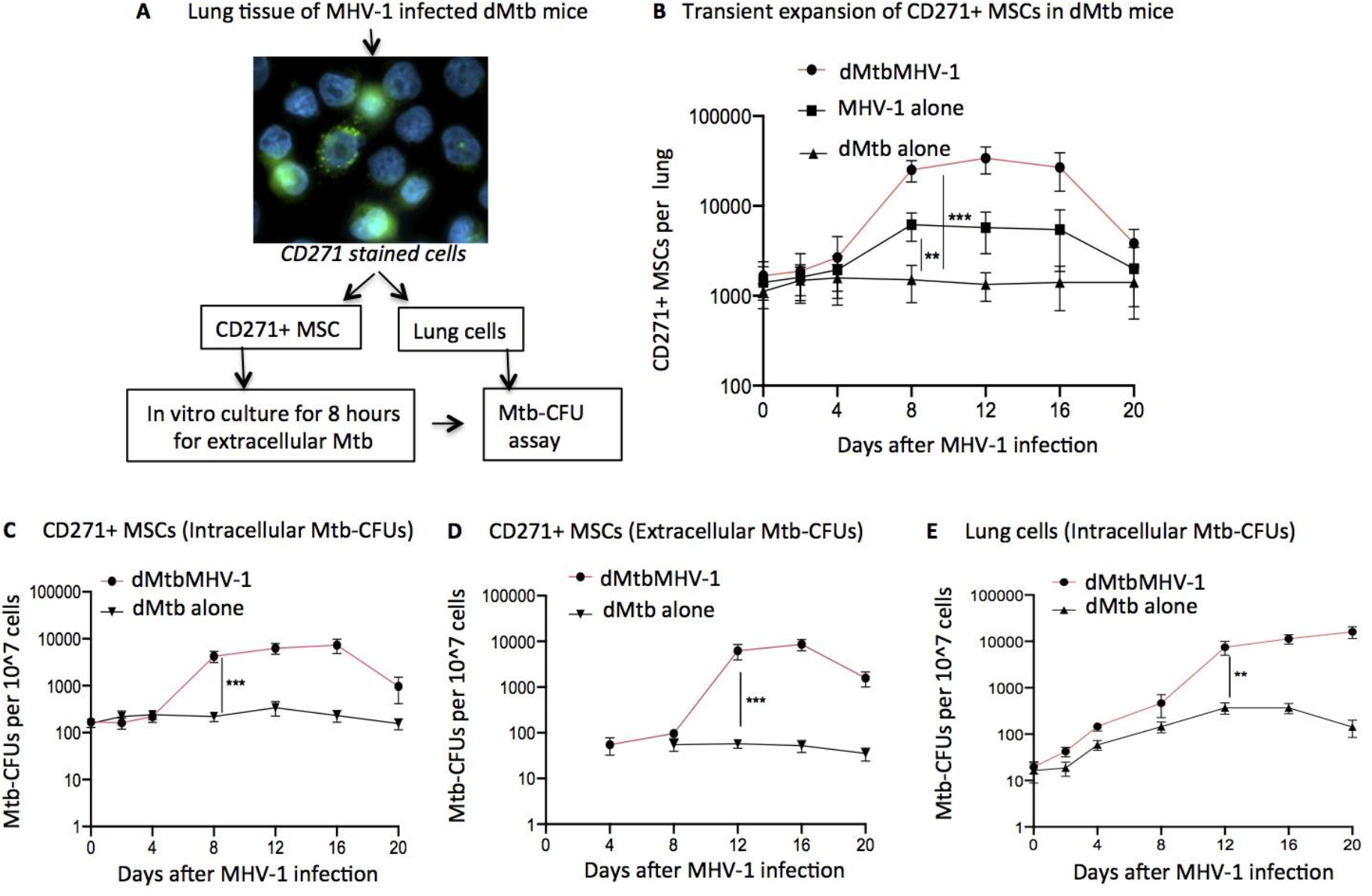
Mtb boost MHV-1 mediated expansion of lung CD271+MSCs. **A**. Schematic of experimental design. Dissociated lung mononuclear cells of MHV-infected mice were obtained at various times post infection. Cells were subjected to immunomagnetic sorting to obtain CD45− and CD271+ mesenchymal stem cells (MSCs) and rest of the lung cells. CD271+ MSCs were then counted by trypan blue and in vitro cultured in serum-free media followed by Mtb-CFU assay for both intra and extracellular Mtb. The leftover lung cells from the immunomagnetic sorting were directly subjected to measure intracellular Mtb-CFU. **B.** Total number of immunomagnetically sorted CD271+/CD45− MSCs in the lungs of MHV-1 infected and non-infected cells. **C&D**. Intracellular and extracellular Mtb-CFUs obtained from CD271+MSCs cultured in vitro for 8-hours. For extracellular Mtb-CFUs, conditioned media was centrifuged, and the pellets were subjected to Mtb-CFU assay. **E.** Intracellular Mtb-CFU/10^7 of non-CD271+ cells (the left over lung mononuclear cells after obtaining CD271+/CD45− MSCs).

### MHV-1 infection activates the altruistic stem cell mediated innate defense mechanism

The MHV-1 infection induced transient expansion of lung CD271+MSCs in both *Mtb* harboring and healthy animal group encouraged us to study the underlying molecular mechanism of expansion. We considered that MHV-1 infection may activate an altruistic stem cell (ASC) based innate defense mechanism that we previously identified in human embryonic stem cells (hESCs) (*22*), and also in mesenchymal stem cells (MSCs) (*18*). Briefly, when a clone of stem cells such as hESCs cell colony is threatened by hypoxia/oxidative stress induced differentiation/apoptosis, a few of these cells reprogram to an “enhanced stemness” phenotype. This phenotype is characterized by distinct gene expression associated with HIF-2alpha stemness pathway (*22*,*29*), and rapid expansion of the reprogrammed cells. These cells continue to maintain enhanced stemness phenotype for the next two weeks, and then activate p53 to undergo differentiation/apoptosis. During the expansion period, these cells exhibit altruistic behaviors by secreting cytoprotective agents such as glutathione that enhance the fitness of the rest of the hES cells. Thus, we identified stem cells with the “enhanced stemness” reprogramming as altruistic Stem Cells (ASCs) (*22*), and described these cells as “intermediate” cell type in Waddington’s epigenetic landscape (*30*) (Figure 3A) (*21*). Indeed, transient cells having ES cell like phenotype has been identified in mice and human, including our findings (*31*) using defined criteria (*32*) (Figure 3B), suggesting the clinical significance of ASCs. We speculate that this ASC reprogramming may function as a putative innate defense system against invading pathogens that threaten the integrity of the stem cells residing in their niches (*32*). Thus, our findings of MHV-1 mediated transient expansion of CD271+MSCs may be part of the ASC defense mechanism against the virus. Indeed, the day-8 CD271+MSCs of MHV-1 infected healthy mice not only showed expansion (Figure 2B), but when subjected to real-time PCR gene expression study (Figure 3C), these cells also exhibited marked alteration of genes associated with enhanced stemness phenotype (HIF-1alpha, HIF-2alpha, Sox2, Oct4, Nanog, ABCG2, MDM2, GCL, p53, and p21) (Figure 3D). We further characterized the stemness of these CD271+MSCs and indeed, they expressed the MSC markers on day 8; CD73, CD74, and Sca-1 whereas CD45 was not expressed (Figure 3D). Demonstration of the increased up-regulation of p53-upstream genes bax, p21 and PUMA, and down-regulation of HIF-2alpha on day-20 (Figure 3E), further confirms the ASC reprogramming. We also noted that the expression of Sca-1 was down regulated in these cells on day 20 (Figure 3E), suggesting differentiation. Interestingly, in the d*Mtb*MHV-1 group, the several ASC reprogramming related gene (Oct4, GCL and ABCG2) expression was increased by 2-3-fold compared to MHV-1 infection alone group (Figure 3D). Thus, these findings suggest that MHV-1 infection led to ASC reprogramming of CD271+MSCs, and this reprogramming was also seen in the d*Mtb* harboring mice, when infected with MHV-1. Activation of p53 upstream genes on day- 20 also indicated the activation of the proposed ASC based innate defense mechanism in the infected CD271+MSCs against MHV-1.

**Figure 3:**
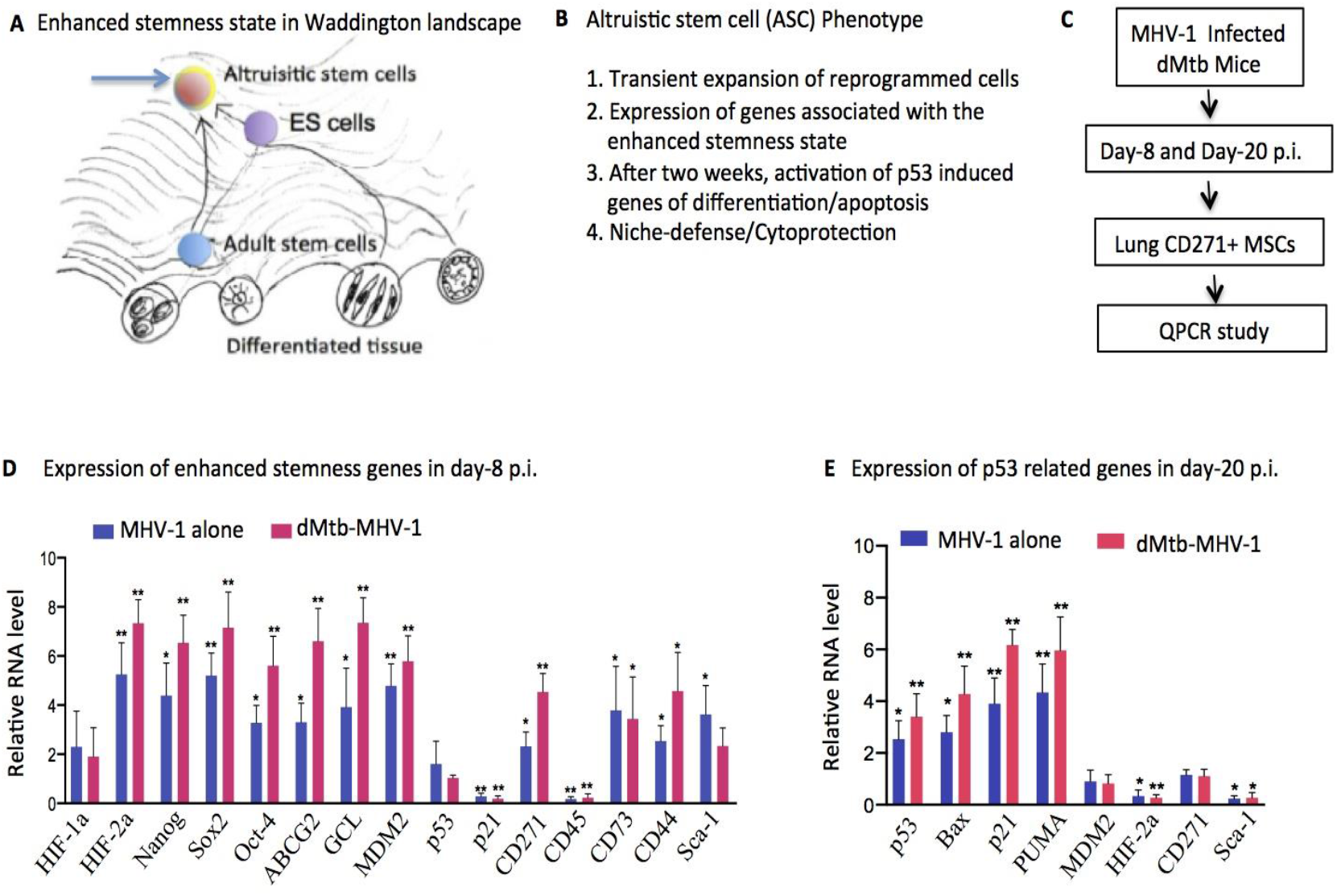
MHV-1 infection reprograms CD271+MSCs to altruistic stem cell phenotype. **A.** Schematic representation to depicting the enhanced stemness phenotype (22) in Waddington’s epigenetic landscape metaphor. Embryonic stem cell (ESCs) and adult stem cells undergo reprogramming to the enhanced stemness state (22) that corresponds to the altruistic stem cell (ASCs) phenotype (21–22,32). ASCs are transient, and therefore, could be considered as the intermediate cell state (21,32), as proposed by Waddington (30). **B**. Phenotypic characteristics of ASCs (32). **C.** Experimental design for qPCR analysis of MHV-1 infected lung CD271+MSCs on day-8 and day-20 post infection. **D**. Histogram showing the induction of enhanced stemness genes in the lung CD271+MSCs of day-8 post-MHV-1 infection. **E**. Histogram showing the induction of p53 related genes of apoptosis/differentiation in lung CD271+ MSCs of day-20 post MHV-1 infection. The qPCR data are expressed as fold-change compared to Mtb-infection alone group. **P< 0.001, ***P< 0.0001, One-Way ANOVA with Dunnett’s post hoc test; error bar represents SEM; n= 4 independent experiments.

We confirmed this ASC based defense against MHV-1 mediated viral load and cellular toxicity in an *in vitro* viral infection assay (Figure 4A). We demonstrated that the conditioned media (CM) of CD271+MSCs on day-8 were capable of exhibiting significant 2-3-fold (p<0.004) cytoprotection of type II lung alveolar epithelial (ATII) cells against MHV-1 (Figure 4B-D). Notably, d*Mtb* presence significantly boosted this ASC based defense against MHV-1 in the *in vitro* viral infection assay, as the viral load was reduced by 3-fold (p<0.0074), while survival of the ATII cells was increased by 2-fold (p<0.001) in the d*Mtb* MHV-1 versus MHV-1 alone group (Figure 4B-C). Thus, these results provide supporting evidence for ASC based defense of lung CD271+MSCs against MHV-1 infection, which could be further enhanced by d*Mtb*.

**Figure 4:**
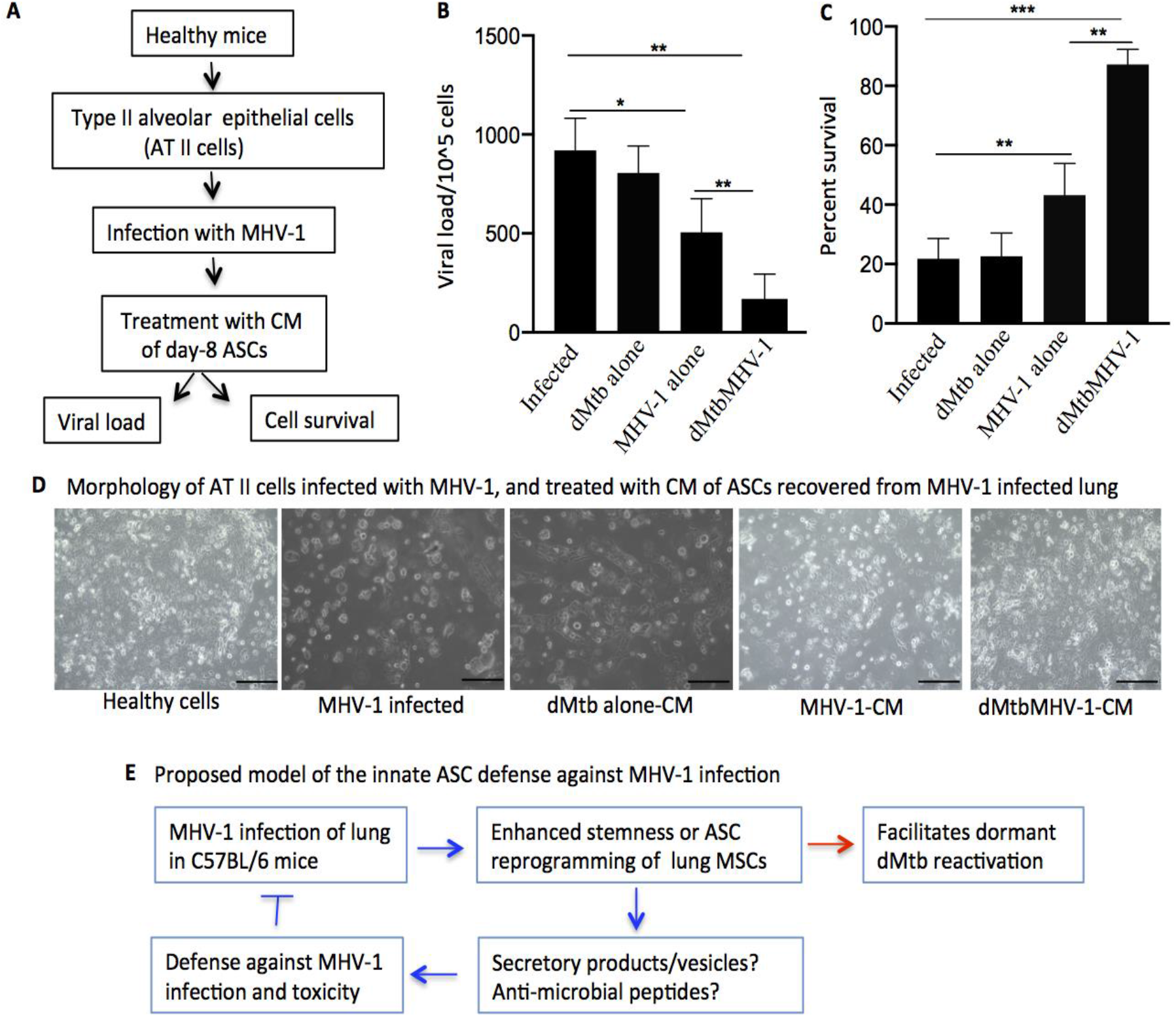
Mtb boost ASC defense against MHV-1 in lung. **A**. Schematic representation of experimental design for ASC defense against MHV-1. **B**. The viral load of type II alveolar epithelial cells (ATII) was obtained after treatment with conditioned media (CM) from lung CD271+MSCs. The CM of lung CD271+MSCs (day-8 post infection) was obtained from all the infected groups. **C**. Cell viability of the treated cells was obtained by trypan blue. The ATII cells not treated with CM served as the control. The CM from CD271+MSCs of dMtbMHV-1 showed the lowest MHV-1-PFU in the ATII cells and highest live cell count. **D**. ATII cells under phase-contrast microscope with magnification 10X after treated with CM of CD271+MSCs obtained from mice lung. **E**. Schematic representation of the niche defense mechanism of MHV-1 infected dMtb harboring mice. ***P< 0.0001, **P< 0.001, *P< 0.05 Student’s t test; Error bar represents SEM; n= 4 independent experiments.

## DISCUSSION

The COVID-19 caused by SARS-CoV-2, has created a global pandemic associated with substantial mortality and morbidity. Notably, a significant number of infected individuals have recovered. However, evidence for a possible host defense or anti-viral mechanism against the virus has not yet been identified. Currently, there are no approved drugs or vaccines to treat COVID-19. Using a mouse coronavirus model, we demonstrate that lung MSCs activate a potential ASC based defense mechanism against the virus and this defense mechanism is further enhanced by d*Mtb* residing inside the lung MSCs. As per recent data from some surveys and reports, it has been seen that countries which do not have a policy of universal *Bacille Calmette Guerin* (BCG) vaccination have experienced higher mortality related to COVID-19 (*33*–*34*). Interestingly, BCG is closely related to *Mtb*, which enhanced ASC based defense mechanism of CD271+MSCs to fight against MHV-1 infection. Thus, our results described here could provide a novel explanation of BCG mediated host defense or anti-viral mechanism against SARS-CoV2 that could be further utilized to design a drug or vaccine against the virus.

In a hostile microenvironment, microorganisms like bacteria activate a unique defense mechanism (*35*), which can be explained as a charitable deed that help the neighbors to survive unfavorable conditions. We have identified a similar altruistic stem cell defense system in hESc, wherein a subset of these cells, (SSEA3+/ABCG2+), acquire an enhanced stemness state that enable these cells to secrete antioxidants and growth factors in the hostile microenvironment of oxidative stress. This altruistic act protects the rest of the population from oxidative stress induced apoptosis and differentiation (*22*). The enhanced stemness state is associated with an increased expression of Oct-4 and Nanog governed by an altered and transient state of the p53/MDM2 oscillation system. After two weeks, the SSEA3+/ABCG2+ ES cell population undergoes apoptosis or differentiation, thus exhibiting an altruistic behavior i.e. enhancing group fitness by sacrificing self-fitness (*22*). In this preliminary study, we have demonstrated an ASC based innate defense response against infection with the mouse coronavirus; MHV-1. The recovered lung CD271+MSCs of MHV-1 infected mice exhibited the ASC phenotype, as shown by their expansion, and expression of genes associated with the HIF-2alpha stemness pathway, followed by the activation of p53 and its downstream genes including p21 and p53 up-regulated modulator of apoptosis (PUMA) on day-20. The activation of genes involved in p53 mediated apoptosis/differentiation was associated with the decrease in lung CD271+MSCs, suggesting that the cells might undergo apoptosis or differentiation, an important phenotypic characteristic of ASCs and ASC based defense mechanism. Notably, further support was obtained for ASC reprogramming *in vitro* by studying the cytoprotective activity of these ASCs in MHV-1 mediated toxicity of lung alveolar epithelial cells. Thus, the conditioned media of the CD271+MSCs recovered from the MHV-1 infected mouse lung, exhibited a direct anti-viral and cytoprotective property on MHV-1 infected type II alveolar epithelial (ATII) cells. Hence, we have identified a unique anti-viral defense system mediated by the innate ASCs. We suggest that ASC defense system may be harnessed to develop novel therapeutic strategies against COVID-19.

Furthermore, our results indicate that *Mtb* infection may boost the MSC mediated ASC defense against MHV-1. Previously, we reported that post-therapy, CD271+BM-MSCs exhibited an enhanced hypoxic phenotype of low CD146 and high HIF-1alpha (*18*), which could modify the stemness state to an altruistic stemness state. We hypothesized that infection and inflammation may activate d*Mtb* intracellular to stem cells by activating ASC based defense mechanism (*18*). Following MHV-1 infection, the non-replicating dormant m18b-harboring CD271+MSCs underwent ASC reprogramming, and permitted m18b replication. There was also a potential transfer of the pathogen to the extracellular space. However, the detailed mechanism of *Mtb* reactivation needs further investigation. Here, we speculate that MHV-1 infection inducing inflammation might trigger the mobilization of d*Mtb* harboring MSCs from BM to lung and cause rapid *Mtb* reactivation in lung, because; (i) studies showed that stem cells mobilize to the area of infection (*20*) (ii) after 6 months of streptomycin starvation, a few Mtb was found in mice lung CD271+MSCs, whereas a greater number of *Mtb* were inside BM-CD271+MSCs (*17*). The remarkable and significant finding at this moment of COVID-19 pandemic is that the *Mtb* reactivation was associated with significant boosting of the ASC mediated anti-viral activity.

Lancet has recently reported that most of the death related to COVID-19 is due to acute respiratory distress syndrome (ARDS), which is characterized by cytokine storm. This will mediate a severe attack by the immune system, as seen in cases of SARS-CoV-1 and MERS-CoV infection, leading to the death of the individual (*36*). Studies support the use of various therapeutic mechanisms of MSCs in the treatment of ARDS. This includes anti-inflammatory effects on host cells, reduction of lung alveolar epithelium permeability, increased rate of alveolar fluid clearance, enhanced phagocytic activity of host mononuclear, direct transfer of mitochondrial DNA and secretion of extracellular vesicles (*54*,*55*). MSCs are immunosuppressive and they release various immune modulators like nitric oxide (NO), prostaglandin (PGE2), interleukin (IL)-6 and IL-10, which can modulate the innate and adaptive immune response (*32*). We speculate that MSC to ASC reprogramming may further increase the secretion of these immunosuppressive cytokines, thus potentially contributing to the reduction of the cytokine storm in COVID-19 subjects treated with MSC therapy.

BCG vaccine has been proposed to work against COVID-19 infection, although the mechanism is not yet known. It is possible that BCG may remain dormant in the lungs, which then reprogram the MSCs to ASCs following COVID-19 infection and thus, could exhibit antiviral defense against the virus. MSCs were already shown to exhibit antiviral activity via extracellular vesicles carrying microRNA (*37*–*39*), and human cathelicidin hCAP-18/ LL-3 (*40*) against influenza, hepatitis C and pneumonia respectively. We have also previously used BCG-MSC interaction to target cancer stem cell (CSCs) (*41*). We speculate that the immunosuppressive property of MSCs could help in neutralizing the severe cytokine storm caused by COVID-19. In SARS-CoV-2 infected humans, BCG might reprogram the MSCs to ASCs. The enhanced stemness state might result in the secretion of factors like immune modulators capable of protecting the cells from damage caused by COVID-19.

The adaptive immune response can play an important role in infection control of MHV-1 in C57BL/6 mice (*25*), however this does not solely account for the resistance displayed by this mouse strain against the viral infection associated morbidity and mortality (*24*). So far in this MHV-1 infection model and immune response, the role of trained immunity has not been evaluated. The trained immunity constitutes innate immune memory where hematopoietic cells may undergo cell intrinsic changes including metabolic and epigenetic modifications to remember pathogenic insult (*42*–*47*). Trained immunity of MSCs has not yet been studied. Whether our findings of ASC mediated innate immunity may include a trained immune component requires further investigation. Such a possibility may provide explanation of BCG-mediated trained immunity against viral infection (*48*–*49*,*56*). Thus, our model of *Mtb* and MHV-1 infection in mice to study the putative ASC defense may be a useful model to study potential MSC based trained immunity against coronavirus, specifically SARS-CoV-2.

A recent report suggest that subjects with TB increases the susceptibility and severity of SARS-CoV-2 (*50*). In contrast, our findings suggest that *Mtb* reactivation reduces the viral load in lung. However our study is based on a mouse model, which needs to be confirmed using human clinical samples. Therefore, we are currently working on COVID-19 subjects to find out *Mtb* reactivation in the lung. We are also studying the presence of circulating ASC in these subjects as potential evidence for SARS-CoV-2 mediated ASC defense mechanism as described previously (*31*).

In summary, we found that MHV-1 infection induces d*Mtb* loaded MSCs to reprogram to ASCs, thus activating the ASC defense mechanism and diminishing the MHV-1 infection. Knowledge of immunity to SARS-CoV2 is still at a preliminary stage. Thus, our study may help in understanding how MSCs inducing ASC defense mechanism will help in combating the viral load in the host; thereby helping in developing a possible cure for COVID-19.

## MATERIALS AND METHODS

### Mycobacterium strain and growth condition

All the necessary experimental procedures were approved and undertaken inside BSC-class II facility in accordance with guidelines of “Institutional Bio-safety Committee” of KaviKrishna Laboratory. Streptomycin-auxotrophic mutant *Mtb* strain18b (gifted by Prof. Stewart T. Cole, Ecole Polytechinque Federale de Lausanne, Lausanne, Switzerland) was cultured in BBL Middlebrook 7H9 broth with glycerol (BD Bioscience, #221832) along with 50 μg/ml of streptomycin sulfate. It was maintained at 37°C and 5% CO_2_ with occasional shaking until the mid-logarithmic phase was reached, OD approximating to 1.

### Development of stem cell mediated mice model of *Mtb* reactivation

All the necessary experimental procedures were undertaken in accordance with approvals of Institutional Animal Ethics Committee, Gauhati University and Institutional Ethics Committee of KaviKrishna Laboratory. The 6-8-week-old C57BL/6 female mice were obtained from National Institution of Nutrition, Hyderabad, India and were maintained in the animal house of Gauhati University at pathogen free condition as previously described (*17*). The mice model of *Mtb* dormancy was developed in 6 months. Briefly, streptomycin-auxotrophic mutant *Mtb* strain18b cell suspension was prepared in phosphate buffered saline (PBS)-Tween 80 (0.05%), sonicated for 15 seconds, and intravenously injected with 2×10^6^ CFUs per mouse. Initially for 3 weeks streptomycin was administered (3mg/mouse in 200 μl of normal saline) daily for establishing infection. Then, mice were observed for 6 months without any streptomycin treatment to establish bacterial dormancy. Following 6 months of streptomycin starvation, lung tissue were dissociated; CD271+/CD45− MSCs and non CD271+ cells were isolated by magnetic sorting as previously explained (18). The magnetically sorted CD271+MSCs and non CD271+ cells were subjected to *Mtb*-CFU assay. Consistent with our previous findings (*17*), a small number of *Mtb*-CFUs (38±10 *Mtb*-CFU/lung) were obtained only in CD271+MSCs. Thus, mice model of *Mtb* dormancy was developed. Then, streptomycin was injected intraperitoneally (3mg/mouse in 200 μl of normal saline) daily for 3 weeks (*17*) with or without MHV-1 infection (*23*,*25*) to develop mice model of reactivation. In a separate experiment, streptomycin treated mice were also treated with immunosuppressive agents dexamethasone (0.08mg/day, 6 times a week) or amino guanidine (2.5% wt/vol in drinking water) to cause d*Mtb* reactivation (*28*). The treated mice were observed for stipulated period of time, and then sacrificed to collect lung tissues. The *Mtb*-CFU of lung was evaluated for *Mtb* reactivation.

### Isolation and culture of CD271+MSCs

The mice lung tissues were dissociated using collagenase/lipase and then, cells were subjected to magnetic sorting to isolate CD271+MSCs as explained previously (*17*–*18*). Briefly, CD45− cells were magnetically sorted using Ter119/CD45-depletion kit (#19771; Stem Cell Technologies, Vancouver, BC). Next, the CD45-cells were subjected to magnetic sorting for isolation of CD271+ cells. First, mouse CD271 antibody (mouse clone ME20.4, catalog number Ab8877; Abcam, Cambridge, MA) was phycoerythrin (PE) conjugated by SiteClick antibody labeling kit (catalog numberS10467; Life Technologies, Grand Island, NY). Next, the EasySep PE sorting kit (catalog number 18554; Stem Cells Technologies) was used to isolate CD271+ cells from the CD45-cell population. For *Mtb*-CFU assay, cells were cultured for 4-8 hours in serum free media without any growth factor. For cytoprotective assay, cells were isolated on day 8 following infection and cultured (1×10^7 cells/ml) in Dulbecco’s modified Eagle’s medium (DMEM)/F12 media without serum for 24 hours to obtain the conditioned media.

### *Mtb*-Colony Forming Unit (CFU) assay

This was carried out as previously described (*17*–*18*). For the whole lung Mtb-CFU, the lung tissue were aseptically removed from sacrificed mice and homogenized in PBS with 0.05% Tween 80, then subjected to CFU assay (*18*). For the Mtb-CFU assay of immunomagnetically sorted cells, cells were isolated by dissociating lung tissue using Collagenase/Lipase (*17*). The pellet was lysed with 1 ml of 0.1% Triton X-100 for 15 min, vortexed for 30 seconds, and serial 10-fold dilution was prepared in Middlebrook 7H9 broth. The diluted whole lung or cell lysate were then separately plated on Middlebrook 7H10 agar plates (BD Bioscience, #295964) along with streptomycin (50μg/ml) for the growth of m18b strain. The agar plates were incubated for 3-4 weeks at 37°C and 5% CO2 and CFUs were counted. CFUs were plotted as the means of log10 CFUs per lung or per 10^7^ cells.

### qPCR assay

To isolate the mRNA from bacterial infected CD271+MSCs, we used the μMACS™ technology (Miltenyi Biotec, Bergisch-Gladbach, Germany) as described previously (*17*). Briefly, the mRNA is isolated in a magnetic column using super paramagnetic Oligo (dT) Microbeads which target the poly RNA tail of mammalian RNA. The mRNA gets attached to the magnetic column, whereas the bacterial and stem cell DNA remained in the passed-through lysate. Then, the mRNA was converted to cDNA in a heated magnetic bar as per manufacture instruction and cDNA was subjected to qPCR analysis using the TaqMan Gene expression assay Biotech, Bergisch-Gladbach, Germany). RNA quantification was done using the ΔΔ Ct method by using the SDS software, version 2.2.1 (*17*). The following TaqMan primers were used: Mouse: CD271 (Mm 00446296_m1), CD45 (Mm 01293575_m1), CD44 (Mm01277163_m1), HIF-2alpha (Mm01236112_m1), ABCG2 (Mm00496364_m1), Oct-4(Mm00658129_g1), Nanog (Mm02019550_s1), Sox-2 (Mm03053810_s1), HIF-1alpha (Mm00468869_m1), p53 (Mm01731290_g1), p21 (Mm01332263_m1), MDM2 (Mm01233136_m1), Glutamate Cysteine Ligase (GCL) (Mm00802655_m1), CD73 (Mm00501915_m1) and GAPDH (Mm99999915_g1).

### MHV-1 infection into mice

Parental virus strain; MHV-1 was obtained from American Type Culture Collection (ATCC) and cultured inside the BSC-class II facility at KaviKrishna Laboratory in accordance with approval of “Institutional Bio-safety Committee” and “Institutional Ethics Committee” of KaviKrishna Laboratory. The animal protocol was approved by the “Institutional Ethics Committee” of KaviKrishna Laboratory and Gauhati University, Guwahati. MHV-1 was propagated on L2 (ATCC HCCL-149) cells, purified by sucrose gradient centrifugation and titrated by end point dilution assay on L2 cells to determine titer of the viral stock as previously described (*51*). Mice (C57BL/6 mice previously infected with the mutant 18b *Mtb* strain followed by 6 months of streptomycin starvation, or control group - 25-26 weeks old healthy mice) were intranasally infected with 5000 PFUs of MHV-1 per mouse (*23*,*25*). Briefly, animals were anesthetized with ketamine intraperitoneally and then 50 μl of MHV-1 in PBS was administered intranasally; this dose is known to cause ARI in this strain of mice (*24*,*26*). Evaluation of lung infection was done by viral load study and measurement of TNF-alpha on day 0 and 6 as described (*23*,*26*).

### Viral plaque forming unit (PFU) assay

This assay was performed as previously described (*51*). In specific time period, animals were euthanized, and the whole lungs were harvested. The lung tissue was homogenized as described (*17*), and the supernatant was stored at −80°C prior to further analysis. One day prior to the formation of the PFU assay, L2 cells were seeded so that the cell density could reach confluence on the day of PFU assay. The serial dilution of lung homogenate derived supernatant was loaded on the L2 monolayer cells and incubated for one hour at 37°C. The inoculum was removed, and it was over layered with methylcellulose media and incubated for 48 hr at 37 °C. The plaques were then counted and the virus titer was calculated using the following formula: titer (PFU/ml) = [(number of plaques/well)/(volume of inoculum/well)] × dilution factor.

### Enzyme Linked Immunosorbent Assay (ELISA) for TNF-- alpha evaluation

This was done by using an Elisa Kit (#MTA00B) of R&D Systems (Minneapolis, MN) as described previously (*22*). Briefly, whole lungs were harvested and homogenized in DMEM/F12 medium supplemented with protease inhibitor mix (Sigma Aldrich). Lung homogenates were centrifuged, and 50 μl of supernatant was added to the wells of the ELISA plate, covered and incubated at room temperature for 2 hours. Each well was aspirated and washed for five times as mentioned in the kit. 100 μl of Mouse TNF-alpha conjugate to horseradish peroxidase was added to each well and covered with an adhesive strip. It was then incubated for 2 hours at room temperature and then aspirated and washed. The plates were developed for 30 minutes using 100 μL of tetramethylbenzidine plus hydroegen peroxide. Plates were read at 450 nm using iMark™ Microplate Absorbance Reader (Biorad, Gurgaon, India).

### In vitro cytoprotection assay against MHV-1 infection

The type II alveolar epithelial (ATII) cells of healthy C57BL/6 mice were first isolated as previously described (*52*) by isolating CD45−, and EPCAM+ cells using immunomagnetic sorting (*53*). The isolated cells were cultured in the fibronectin coated 6-well plates (2×10^5 cells/well) using DMEM/F12 and 10% FBS with penicillin/streptomycin antibiotics. The cells were infected with MHV-1 with a MOI of 1:5 as described (*52*) and then treated for 48 hours with the CM obtained from the lung CD271+MSCs collected from the three groups (d*Mtb* alone, MHV-1 alone and the d*Mtb*MHV-1 group). The conditioned media (CM) was collected from *in vivo* CD271+ MSCs by a previously described method (*22*), where the isolated CD271+MSCs (1×10^7 cells/ml) were cultured *in vitro* for 24 hours, and then CM was filtered through a 0.2 micron filter. The filtered CM was added to ATII cells. After 48 hours of growth, cytoprotection was measured by the reduction in viral load (viral plaque assay), and cell survival (trypan blue assay). Trypan blue assay was done as previously described (*22*).

### Statistics

Statistical analysis was carried out using GraphPad Prism version 4.0 (Hearne Scientific Software, Chicago, IL, USA). Student’s t test was used for comparisons with Newman-Keul post hoc test. The gene expression was analyzed using One-Way ANOVA with Dunnett’s post-hoc test. Data are expressed as means ± SEM; *P < 0.05, **P < 0.001, ***P < 0.0001.

## ACKNOWLEDGEMENT

We thank the members of Department of Stem Cell and Infectious Diseases, KaviKrishna Laboratory, Guwahati Biotech Park, Indian Institute of Technology, Guwahati, India, Department of Stem Cell and Infection, Thoreau Lab for Global Health, University of Massachusetts, Lowell, US, Department of Bioengineering and Technology, Gauhati University, Guwahati, India. This research project was funded by grants from the, Laurel Foundation (BD), KaviKrishna Foundation, Assam, India (LP, SG, SB, JT and SS), Department of Biotechnology (DBT) India (BT/PR22952/NER/95/572/2017) (BD) and KaviKrishna USA Foundation (BD and BP).

## AUTHOR CONTRIBUTIONS

BD initiated and designed the study. BD, LP, SG, BP, SB, SS and JT performed the *in vitro* and *in vivo* experiments. BD, LP, SG and BP analyzed data. HY provided materials and reagents. DB provided access to animal facility, and took care of ethical permissions. BD, LP and SG wrote the paper.

## REFERENCES

1. Walaza S, Tempia S, Dawood H, Variava E, Wolter N, Dreyer A, Moyes J, Von Mollendorf C, McMorrow M, Von Gottberg A, Haffejee S, Venter M, Treurnicht FK, Hellferscee O, Martinson NA, Ismail N, Cohen C. The Impact of Influenza and Tuberculosis Interaction on Mortality Among Individuals Aged >/=15 Years Hospitalized With Severe Respiratory Illness in South Africa, 2010-2016. Open Forum Infect Dis.6(3):ofz020. (2019).

2. Oei W, Nishiura H. The relationship between tuberculosis and influenza death during the influenza (H1N1) pandemic from 1918-19. Comput Math Methods Med.2012:124861. (2012).

3. Walaza S, Cohen C, Tempia S, Moyes J, Nguweneza A, Madhi SA, McMorrow M, Cohen AL. Influenza and tuberculosis co-infection: A systematic review. Influenza Other Respir Viruses.14(1):77–91. (2020).

4. Low JG, Lee CC, Leo YS. Severe acute respiratory syndrome and pulmonary tuberculosis. Clin Infect Dis.38(12):e123–5. (2004).

5. Alfaraj SH, Al-Tawfiq JA, Altuwaijri TA, Memish ZA. Middle East Respiratory Syndrome Coronavirus and Pulmonary Tuberculosis Coinfection: Implications for Infection Control. Intervirology.60(1-2):53–5. (2017).

6. Redford PS, Mayer-Barber KD, McNab FW, Stavropoulos E, Wack A, Sher A, O’Garra A. Influenza A virus impairs control of Mycobacterium tuberculosis coinfection through a type I interferon receptor-dependent pathway. J Infect Dis.209(2):270–4. (2014).

7. de Paus RA, van Crevel R, van Beek R, Sahiratmadja E, Alisjahbana B, Marzuki S, Rimmelzwaan GF, van Dissel JT, Ottenhoff TH, van de Vosse E. The influence of influenza virus infections on the development of tuberculosis. Tuberculosis (Edinb).93(3):338–42. (2013).

8. Mamelund SE, Dimka J. Tuberculosis as a Risk Factor for 1918 Influenza Pandemic Outcomes. Trop Med Infect Dis.4(2). (2019).

9. Walaza S, Tempia S, Dawood H, Variava E, Moyes J, Cohen AL, Wolter N, Groome M, von Mollendorf C, Kahn K, Pretorius M, Venter M, Madhi SA, Cohen C. Influenza virus infection is associated with increased risk of death amongst patients hospitalized with confirmed pulmonary tuberculosis in South Africa, 2010-2011. BMC Infect Dis.15:26. (2015).

10. Pandemic influenza A (H1N1) 2009: considerations for tuberculosis care services, Paul Nunn (Coordinator) and Dennis Falzon (Medical Officer), Stop TB Department, World Health Organisation, Geneva. 12 November, (2009).

11. Park Y, Chin BS, Han SH, Yun Y, Kim YJ, Choi JY, Kim CO, Song YG, Kim JM. Pandemic Influenza (H1N1) and Mycobacterium tuberculosis Co-infection. Tuberc Respir Dis (Seoul).76(2):84–7. (2014).

12. Volkert M, Pierce C, Horsfall FL, Dubos RJ. THE ENHANCING EFFECT OF CONCURRENT INFECTION WITH PNEUMOTROPIC VIRUSES ON PULMONARY TUBERCULOSIS IN MICE. J Exp Med.86(3):203–14. (1947).

13. Co DO, Hogan LH, Karman J, Heninger E, Vang S, Wells K, Kawaoka Y, Sandor M. Interactions between T cells responding to concurrent mycobacterial and influenza infections. J Immunol.177(12):8456–65. (2006).

14. Okabayashi T, Kariwa H, Yokota S, Iki S, Indoh T, Yokosawa N, Takashima I, Tsutsumi H, Fujii N. Cytokine regulation in SARS coronavirus infection compared to other respiratory virus infections. J Med Virol.78(4):417–24. (2006).

15. Kaufmann SH, Dorhoi A. Inflammation in tuberculosis: interactions, imbalances and interventions. Curr Opin Immunol.25(4):441–9. (2013).

16. Beamer G, Major S, Das B, Campos-Neto A. Bone marrow mesenchymal stem cells provide an antibiotic-protective niche for persistent viable Mycobacterium tuberculosis that survive antibiotic treatment. Am J Pathol.184(12):3170–5. (2014).

17. Das B, Kashino SS, Pulu I, Kalita D, Swami V, Yeger H, Felsher DW, Campos-Neto A. CD271(+) bone marrow mesenchymal stem cells may provide a niche for dormant Mycobacterium tuberculosis. Sci Transl Med.5(170):170ra13. (2013).

18. Garhyan J, Bhuyan S, Pulu I, Kalita D, Das B, Bhatnagar R. Preclinical and Clinical Evidence of Mycobacterium tuberculosis Persistence in the Hypoxic Niche of Bone Marrow Mesenchymal Stem Cells after Therapy. Am J Pathol.185(7):1924–34. (2015).

19. Jones E, McGonagle D. Human bone marrow mesenchymal stem cells in vivo. Rheumatology (Oxford).47(2):126–31. (2008).

20. Spaeth EL, Kidd S, Marini FC. Tracking inflammation-induced mobilization of mesenchymal stem cells. Methods Mol Biol.904:173–90. (2012).

21. Das B. Altruistic stem cells and cancer stem cells.p. 89–106 In: Rajasekhar VK, editor. Cancer Stem Cells. Hoboken, NJ: John Wiley & Sons. (2014).

22. Das B, Bayat-Mokhtari R, Tsui M, Lotfi S, Tsuchida R, Felsher DW, Yeger H. HIF-2alpha suppresses p53 to enhance the stemness and regenerative potential of human embryonic stem cells. Stem Cells.30(8):1685–95. (2012).

23. De Albuquerque N, Baig E, Ma X, Zhang J, He W, Rowe A, Habal M, Liu M, Shalev I, Downey GP, Gorczynski R, Butany J, Leibowitz J, Weiss SR, McGilvray ID, Phillips MJ, Fish EN, Levy GA. Murine hepatitis virus strain 1 produces a clinically relevant model of severe acute respiratory syndrome in A/J mice. J Virol.80(21):10382–94. (2006).

24. Khanolkar A, Fulton RB, Epping LL, Pham NL, Tifrea D, Varga SM, Harty JT. T cell epitope specificity and pathogenesis of mouse hepatitis virus-1-induced disease in susceptible and resistant hosts. J Immunol.185(2):1132–41. (2010).

25. Khanolkar A, Hartwig SM, Haag BA, Meyerholz DK, Harty JT, Varga SM. Toll-like receptor 4 deficiency increases disease and mortality after mouse hepatitis virus type 1 infection of susceptible C3H mice. J Virol.83(17):8946–56. (2009).

26. Khanolkar A, Hartwig SM, Haag BA, Meyerholz DK, Epping LL, Haring JS, Varga SM, Harty JT. Protective and pathologic roles of the immune response to mouse hepatitis virus type 1: implications for severe acute respiratory syndrome. J Virol.83(18):9258–72. (2009).

27. Kashino SS, Ovendale P, Izzo A, Campos-Neto A. Unique model of dormant infection for tuberculosis vaccine development. Clin Vaccine Immunol.13(9):1014–21. (2006).

28. Scanga CA, Mohan VP, Joseph H, Yu K, Chan J, Flynn JL. Reactivation of latent tuberculosis: variations on the Cornell murine model. Infect Immun.67(9):4531–8. (1999).

29. Das B, Pal B, Bhuyan R, Li H, Sarma A, Gayan S, Talukdar J, Sandhya S, Bhuyan S, Gogoi G, Gouw AM, Baishya D, Gotlib JR, Kataki AC, Felsher DW. MYC Regulates the HIF2alpha Stemness Pathway via Nanog and Sox2 to Maintain Self-Renewal in Cancer Stem Cells versus Non-Stem Cancer Cells. Cancer Res.79(16):4015–25. (2019).

30. Waddington CH. Canalization of Development and the Inheritance of Aquired Characters.. Nature.150(3811):563–5. (1942).

31. Talukdar J, Bhuyan R, Garhyan J, Pal B, Sandhya S, Gayan S, Sarma A, Mokhtari R B, Li H, Phukan J, Tasabehji W, Bhuyan S, Kataki A C, Tsuchida R, Yeger H, Baishya D, Das B. Migratory cancer side population cells induces stem cell altruism in bone marrow mesenchymal stem cells to resist therapy, and enhance tumorigenic potential of non-tumorigenic cells. In: Proceedings of the 107th Annual Meeting of the American Association for Cancer Research, New Orleans, LA Philadelphia (PA):AACR;Cancer Res 2016;76(14 Suppl):Abstract nr 920. (2016).

32. Pal B, Das B. In vitro Culture of Naive Human Bone Marrow Mesenchymal Stem Cells: A Stemness Based Approach. Front Cell Dev Biol.5:69. (2017).

33. Miller A RMJ, Fasciglione K, Roumenova V, Li Y, Otazu G H. 2020. Correlation between universal BCG vaccination policy and reduced morbidity and mortality for COVID-19: an epidemiological study. 10.1101/2020.03.24.20042937: Available from: https://www.medrxiv.org/content/10.1101/2020.03.24.20042937v1.

34. Redelman-Sidi G. Could BCG be used to protect against COVID-19? Nature Reviews Urology. (2020).

35. Lee HH, Molla MN, Cantor CR, Collins JJ. Bacterial charity work leads to population-wide resistance. Nature.467(7311):82–5. (2010).

36. Huang C, Wang Y, Li X, Ren L, Zhao J, Hu Y, Zhang L, Fan G, Xu J, Gu X, Cheng Z, Yu T, Xia J, Wei Y, Wu W, Xie X, Yin W, Li H, Liu M, Xiao Y, Gao H, Guo L, Xie J, Wang G, Jiang R, Gao Z, Jin Q, Wang J, Cao B. Clinical features of patients infected with 2019 novel coronavirus in Wuhan, China. Lancet.395(10223):497–506. (2020).

37. Qian X, Xu C, Fang S, Zhao P, Wang Y, Liu H, Yuan W, Qi Z. Exosomal MicroRNAs Derived From Umbilical Mesenchymal Stem Cells Inhibit Hepatitis C Virus Infection. Stem Cells Transl Med.5(9):1190–203. (2016).

38. Khatri M, Richardson LA, Meulia T. Mesenchymal stem cell-derived extracellular vesicles attenuate influenza virus-induced acute lung injury in a pig model. Stem Cell Res Ther.9(1):17. (2018).

39. Yang K, Wang J, Wu M, Li M, Wang Y, Huang X. Mesenchymal stem cells detect and defend against gammaherpesvirus infection via the cGAS-STING pathway. Sci Rep.5:7820. (2015).

40. Krasnodembskaya A, Song Y, Fang X, Gupta N, Serikov V, Lee JW, Matthay MA. Antibacterial effect of human mesenchymal stem cells is mediated in part from secretion of the antimicrobial peptide LL-37. Stem Cells.28(12):2229–38. (2010).

41. Pal B, Bhuyan S, Garhyan J, Li H, Bhuyan R, Yeger H, Das B. Targeting oral cancer stem cells in the hypoxic niche by BCG infected mesenchymal stem cells. In: Proceedings of the American Association for Cancer Research Annual Meeting, Washington, DC Philadelphia (PA),:AACR; Cancer Res 2017;77(13 Suppl):Abstract nr 3903. (2017).

42. Song WM, Colonna M. Immune Training Unlocks Innate Potential. Cell.172(1-2):3–5. (2018).

43. Netea MG, Quintin J, van der Meer JW. Trained immunity: a memory for innate host defense. Cell Host Microbe.9(5):355–61. (2011).

44. Gourbal B, Pinaud S, Beckers GJM, Van Der Meer JWM, Conrath U, Netea MG. Innate immune memory: An evolutionary perspective. Immunol Rev.283(1):21–40. (2018).

45. Netea MG. Training innate immunity: the changing concept of immunological memory in innate host defence. Eur J Clin Invest.43(8):881–4. (2013).

46. Bekkering S, Arts RJW, Novakovic B, Kourtzelis I, van der Heijden C, Li Y, Popa CD, Ter Horst R, van Tuijl J, Netea-Maier RT, van de Veerdonk FL, Chavakis T, Joosten LAB, van der Meer JWM, Stunnenberg H, Riksen NP, Netea MG. Metabolic Induction of Trained Immunity through the Mevalonate Pathway. Cell.172(1-2):135–46 e9. (2018).

47. Mitroulis I, Ruppova K, Wang B, Chen LS, Grzybek M, Grinenko T, Eugster A, Troullinaki M, Palladini A, Kourtzelis I, Chatzigeorgiou A, Schlitzer A, Beyer M, Joosten LAB, Isermann B, Lesche M, Petzold A, Simons K, Henry I, Dahl A, Schultze JL, Wielockx B, Zamboni N, Mirtschink P, Coskun U, Hajishengallis G, Netea MG, Chavakis T. Modulation of Myelopoiesis Progenitors Is an Integral Component of Trained Immunity. Cell.172(1-2):147–61 e12. (2018).

48. Arts RJW, Moorlag S, Novakovic B, Li Y, Wang SY, Oosting M, Kumar V, Xavier RJ, Wijmenga C, Joosten LAB, Reusken C, Benn CS, Aaby P, Koopmans MP, Stunnenberg HG, van Crevel R, Netea MG. BCG Vaccination Protects against Experimental Viral Infection in Humans through the Induction of Cytokines Associated with Trained Immunity. Cell Host Microbe.23(1):89–100 e5. (2018).

49. Christ A, Gunther P, Lauterbach MAR, Duewell P, Biswas D, Pelka K, Scholz CJ, Oosting M, Haendler K, Bassler K, Klee K, Schulte-Schrepping J, Ulas T, Moorlag S, Kumar V, Park MH, Joosten LAB, Groh LA, Riksen NP, Espevik T, Schlitzer A, Li Y, Fitzgerald ML, Netea MG, Schultze JL, Latz E. Western Diet Triggers NLRP3-Dependent Innate Immune Reprogramming. Cell.172(1-2):162–75 e14. (2018).

50. Liu Y, Bi L, Chen Y, Wang Y, Fleming J, Yu Y, Gu Y, Liu C, Fan L, Wang X, Cheng M. Active or latent tuberculosis increases susceptibility to COVID-19 and disease severity. medRxiv:2020.03.10.20033795. (2020).

51. Leibowitz J, Kaufman G, Liu P. Coronaviruses: propagation, quantification, storage, and construction of recombinant mouse hepatitis virus. Curr Protoc Microbiol.Chapter 15:Unit 15E 1. (2011).

52. Kebaabetswe LP, Haick AK, Miura TA. Differentiated phenotypes of primary murine alveolar epithelial cells and their susceptibility to infection by respiratory viruses. Virus Res.175(2):110–9. (2013).

53. Pal B, Bhuyan S, Baishya D, Das B. Oral cancer stem cells modulate Fusobacterium nucleatumto acquire the capability to induce tumor stemness switch. In: Proceedings of the American Association for Cancer Research Annual Meeting, Chicago, IL Philadelphia (PA):AACR; Cancer Res 2018;78(13 Suppl):Abstract nr 3064. (2018).

54. Huppert LA, Liu KD, Matthay MA. Therapeutic potential of mesenchymal stromal ells in the treatment of ARDS. Transfusion 59: 869–875. (2019).

55. Park J, Kim S, Lim H, Liu A, Hu S, Lee J, Zhuo H, Hao Q, Matthay MA, and Lee JW. Therapeutic Effects of Human Mesenchymal Stem Cell Microvesicles in an Ex Vivo Perfused Human Lung Injured with Severe E.coli Pneumonia. Thorax 4(1): 43–50. (2019).

56. O’Neill LAJ and Netea MG. BCG-induced trained immunity: can it offer protection against COVID-19? Nat Rev Immunol. 1–3. (2020).

